# Morphological and phenotypic characterization of adipose-derived mesenchymal stem cells isolated from locally adapted Indonesian goat breed

**DOI:** 10.64898/2026.08.23.746531

**Authors:** Teguh Budipitojo, Irma Padeta, Medania Purwaningrum, Vista Budiariati, Nopadon Pirarat

## Abstract

Adipose-derived mesenchymal stem cells (gAD-MSCs) are promising candidates for veterinary regenerative medicine, yet the characterization of gAD-MSCs from locally adapted Indonesian goat breeds remains limited. This study aimed to isolate and characterize gAD-MSCs from Peranakan Ettawa (PE) goats using tissue explant culture. Subcutaneous adipose tissue was collected from the base of the tail of healthy PE goats (n=3). Primary cell outgrowth from explants was observed by Day 5, displaying characteristic fibroblast-like, spindle-shaped morphology and strong plastic adherence. Serial passaging to Passage 3 (P3) yielded a morphologically stable, homogeneous cell population. Assessment of cellular metabolic activity via the resazurin assay demonstrated sustained cell viability and a statistically significant increase in metabolic activity between Day 3 and Day 5 (*p* < 0.05). Furthermore, functional clonogenic capacity, evaluated using the colony-forming unit (CFU) assay, showed continuous temporal expansion of colonies over 14 days, yielding an average of 52.0 ± 4.1 colonies per dish. These findings confirm that expanded gAD-MSCs P3from PE goats maintain characteristic mesenchymal morphology, sustained metabolic activity, and clonogenic capacity. This work provides a baseline cellular profile of PE goat gAD-MSCs, supporting their potential use in veterinary regenerative medicine and tissue engineering.

## INTRODUCTION

Mesenchymal stem cells (MSCs) are multipotent cells that have attracted considerable interest in regenerative medicine because of their capacity for self-renewal, multilineage differentiation, and secretion of bioactive factors involved in tissue repair and immunomodulation (1, 2). Mesenchymal stem cells (MSCs) can be obtained from various tissues, including bone marrow, adipose tissue, umbilical cord, and placenta. Among these sources, adipose tissue is particularly attractive because it is relatively abundant and accessible, allowing the isolation of adipose-derived MSCs (AD-MSCs) via minimally invasive procedures and offering substantial expansion potential (3).

Goats (*Capra hircus*) are important livestock species and have also been used as large-animal models in regenerative medicine and tissue engineering research. However, compared with human MSCs and MSCs from commonly studied laboratory or companion animals, goat MSCs remain comparatively less well characterized. Previous studies demonstrated that goat MSCs isolated from adipose tissue and bone marrow exhibit osteogenic, adipogenic, and chondrogenic differentiation, while differences in differentiation capacity and other cellular characteristics can occur according to tissue source and culture conditions (4, 5). More recently, goat AD-MSCs have been shown to maintain fibroblast-like morphology, clonogenic capacity, proliferation, mesenchymal-associated surface-marker expression, and multilineage differentiation under appropriate culture conditions (6).

Characterization of MSCs generally includes assessment of cell morphology, adherence to plastic, surface-marker expression, proliferative or clonogenic capacity, and multilineage differentiation. The International Society for Cellular Therapy (ISCT) proposed minimal criteria for defining human MSCs, including plastic adherence, expression of characteristic mesenchymal markers, absence of hematopoietic markers, and differentiation into adipogenic, chondrogenic, and osteogenic lineages (1). However, the expression of individual surface markers may vary among species, and trilineage differentiation has been suggested as a more consistent functional characteristic for identifying MSCs across different animal species (4).

In Indonesia, Peranakan Ettawa (PE) local goats represent an important small-ruminant resource, while their cells may provide a valuable source for veterinary regenerative medicine and biological research. Nevertheless, the comprehensive characterization of AD-MSCs derived from local goats remains limited. The previous research framework underlying this study identified the need to characterize MSCs from local small ruminants based on morphology, proliferation, phenotype, and multilineage differentiation, with adipose tissue representing one of the principal tissue sources. Therefore, this study aimed to isolate AD-MSCs from local goats and analyze their phenotypic characteristics. The findings are expected to provide a baseline characterization of local goat AD-MSCs and support their potential use in subsequent veterinary regenerative medicine and tissue-engineering studies.

## MATERIALS AND METHODS

### Adipose tissue collection and adipose-derived mesenchymal stem cells isolation

Adipose tissue was obtained from healthy Peranakan Ettawa (PE) goat, a locally adapted Indonesian goat breed (n=3) aged 1-1.5 years old, weighing 4-5 kg, maintained at the Animal Health Education and Training Unit (UP2KH), Faculty of Veterinary Medicine, Universitas Gadjah Mada. Adipose tissue was collected from the subcutaneous adipose tissue at the base of the tail using an aseptic procedure. The collected tissue was immediately transferred into a sterile transport medium. All animal procedures were performed following approval from the animal ethics committee under protocol number 03/EC-FKH/Int./2026.

Adipose-derived MSCs were isolated through the tissue explant technique. Briefly, adipose tissue was transferred and repeatedly washed with sterile phosphate-buffered saline (PBS, Procell System, China) supplemented with an antibiotic-antimycotic solution to remove blood and other tissue contaminants. Visible connective tissue and non-adipose components were removed where necessary. The adipose tissue was then minced into small fragments. The minced tissue was transferred into 60-mm TC dish (Labselect, China) containing Dulbecco’s Modified Eagle Medium (DMEM, Procell System) supplemented with 10% Fetal Bovine Serum (FBS, BioWhittaker, Switzerland), 1% GlutaMAX (ThermoFisher, USA), and 1% Antibiotic-Antimycotic (ABAM, ThermoFisher). Tissues were maintained at 37°C in a humidified atmosphere containing 5% CO□.

Cells derived from explant culture were maintained until it reached approximately 80% confluence, and they were detached using 0.25% Trypsin-EDTA (ThermoFisher) and subcultured int new TC dishes. Cells were expanded until passage 3-4 (P3-4) for subsequent characterization. Expanded culture was also preserved as a cryostock of gAD-MSCs in liquid nitrogen.

### Cell proliferation assay

Primary characterization of isolated gAD-MSCs included morphological observation under a 10X and 20X plan phase-contrast objective using Oxion Inverso Microscope (EUromex, Netherlands). Cell proliferation of gAD-MSCs at P3 was assessed using the resazurin assay in a 96-well plate (Lab Select). Initial seeding cell density was 2 × 10^3^/well, maintained in complete DMEM at 37°C in a humidified atmosphere containing 5% CO . Assessment was employed on days 5, 7, and 9 after incubation with 0.015 mg/mL resazurin sodium salt (Sigma, USA). After 3 h of incubation, absorbance was measured using Accuris SmartReader UV-Vis (Benchmark Scientific, China) at 570 and 600 nm. Resazurin reduction was interpreted as an increase in metabolically active cells, indicating the number of viable cells.

### Colony-forming unit (CFU) assay

The self-renewal and survival capacity of gAD-MSCs were investigated using the colony-forming unit assay. Initial cell seeding density was 3 × 10^3^ cells per 60-mm TC dish, maintained in complete DMEM at 37°C in a humidified atmosphere containing 5% CO for 14 days. Following incubation, colonies were stained with crystal violet solution (Merck Millipore, USA) and counted for analysis. A gAD-MSCs colony was defined as a cluster containing ≥50 cells. The CFU assay was employed gradually on Day 5, 10, and 14.

### Statistical analysis

The morphology and colony of gAD-MSCs were observed using ImageJ Software (https://imagej.net/ij/). The visible field of view (FOV) was captured and calibrated using the gridded area of the Neubauer Hemocytometer. Resazurin result and colony number were. The results of this study were interpreted using a dot plot in GraphPad Prism 9.0 (GraphPad Software, Inc., CA). The illustrated data were analyzed using IBM SPSS Statistics 29 (IBM Corporation, USA). The Kruskal-Wallis test was employed to compare significant differences among more than two experimental groups, and statistical significance was interpreted by *p* <0.05.

## RESULT

### Adherent fibroblast-like morphology supports the initial characterization of PE gAD-MSCs

Goat adipose-derived mesenchymal stem cells (gAD-MSCs) were successfully isolated from adipose tissue collected from the base of the tail of Peranakan Ettawa (PE) goats, using the explant culture technique. Cell migration from the tissue explants was first observed on Day 5 of culture, with a small number of adherent cells displaying an elongated spindle-shaped morphology. By Day 11, the outgrowing cell population had markedly increased, forming a denser layer of cells surrounding the explanted tissue. The isolated cells exhibited a fibroblast-like morphology characterized by elongated cell bodies with centrally positioned nuclei and strong adherence to the culture surface (Figure 1). Furthermore, cell proliferation occurred locally around the explants, resulting in progressive expansion of the adherent cell population. These morphological characteristics are consistent with the typical features of mesenchymal stem cells during primary culture.

**Figure 1.**
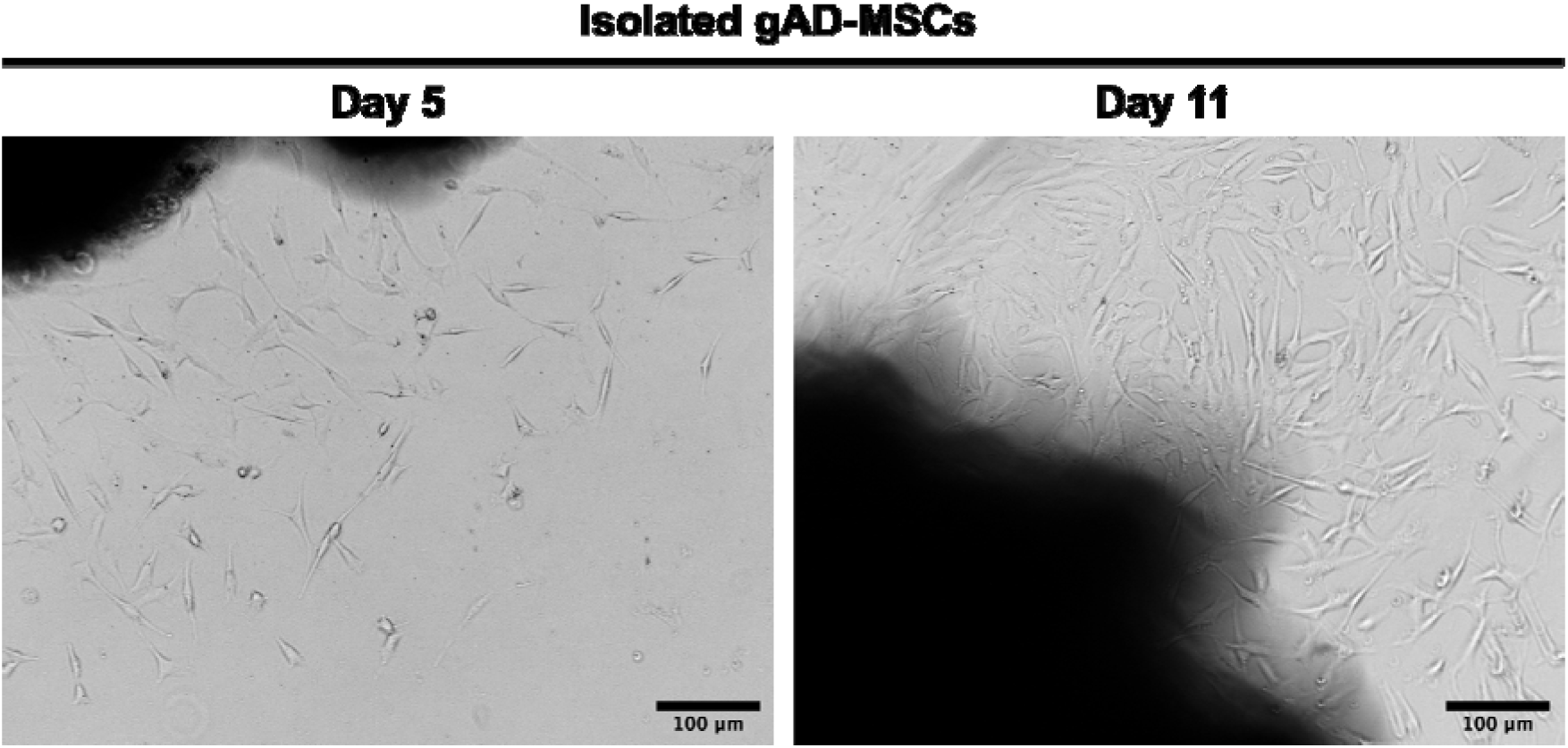
Representative micrographs of gAD-MSC isolation from tail-base adipose tissue explants. Cells migrated from the adipose tissue explants and attached to the culture surface during primary culture. Sparse spindle-shaped cells were observed on Day 5, whereas a greater number of cells with typical mesenchymal morphology were present on Day 11. The cells displayed fibroblast-like morphology and local proliferation surrounding the explanted tissue. Scale bar = 100 μm.

Expansion of gAD-MSCs to passage 3 resulted in the establishment of a morphologically stable and relatively homogeneous cell population (Figure 2). The cultured cells consistently exhibited strong adherence to the plastic culture surface and retained the characteristic fibroblast-like morphology of mesenchymal stem cells, including elongated spindle-shaped cell bodies with centrally positioned nuclei. Serial passaging progressively reduced the presence of non-mesenchymal cellular phenotypes, yielding a population with minimal morphological variation and high uniformity. The preservation of these characteristic morphological features during expansion indicates effective propagation of the mesenchymal cell population and provides morphological evidence supporting the use of P3 cells for subsequent functional and phenotypic characterization.

**Figure 2.**
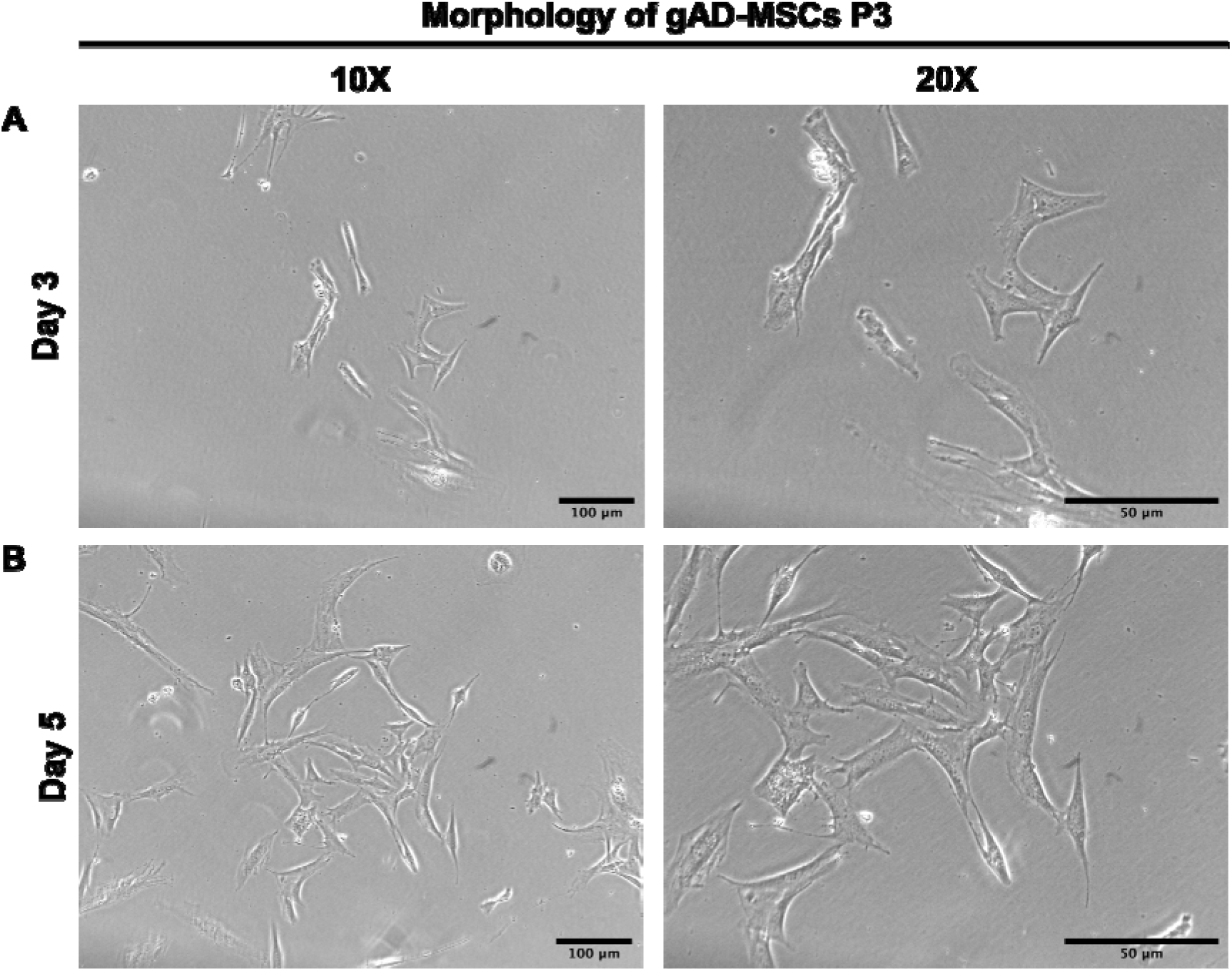
Morphological characteristics and expansion of gAD-MSCs at Passage 3. Expanded gAD-MSCs maintained stable adherence and exhibited a uniform fibroblast-like morphology characterized by elongated spindle-shaped cells with centrally positioned nuclei. The progressive reduction in morphological heterogeneity during culture expansion resulted in a relatively homogeneous cell population, consistent with the expected morphology of mesenchymal stem cells. Scale bars = 100 μm and 50 μm.

### Increasing resazurin reduction indicates sustained metabolic activity of PE gAD-MSCs

The proliferative behavior of gAD-MSCs passage 3 was assessed using the resazurin assay after seeding 2 × 10^3^ cells. Cell viability increased progressively with culture duration, as evidenced by a gradual reduction in resazurin assay (Figure 3). A significant increase was detected between Day 3 and Day 5 (*p*<0.05), indicating a marked increase in metabolic activity during this culture period. The continued evaluation of the resazurin signal at subsequent time points was consistent with progressive accumulation of viable, metabolically active cells. These results demonstrate that passage 3 gAD-MScs maintained viability following initial seeding and exhibited sustained growth-associated metabolic activity under the established culture conditions.

**Figure 3.**
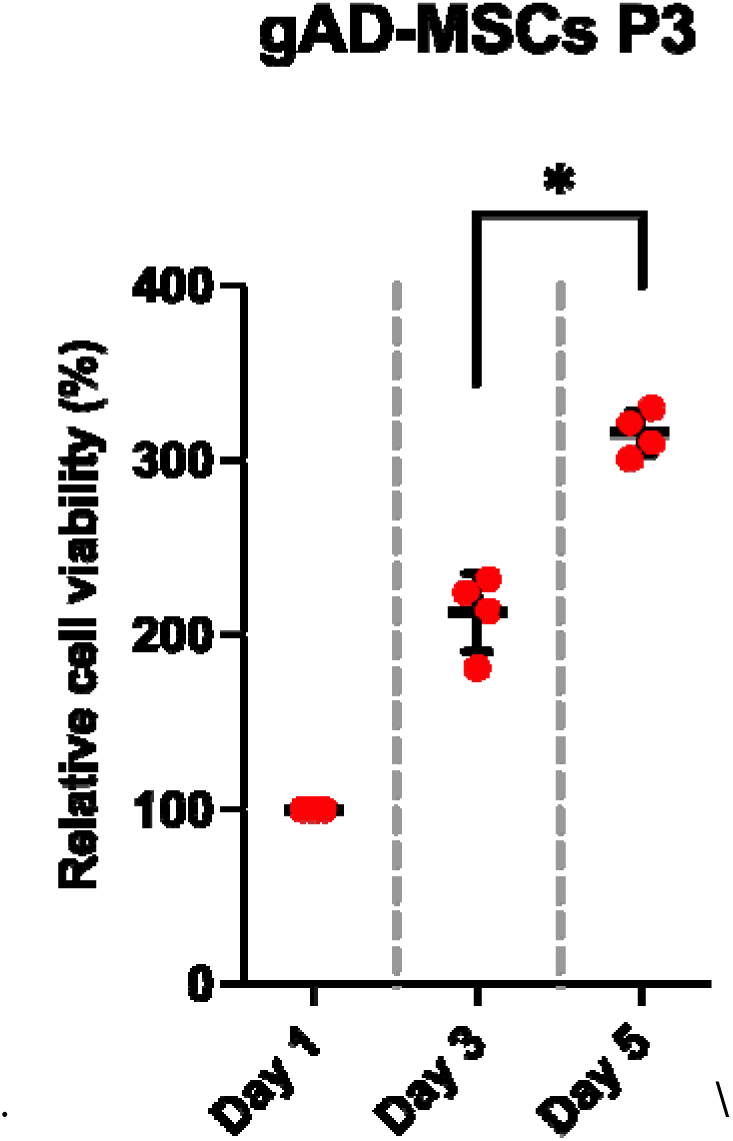
Relative cell viability of gAD-MSCs during in vitro culture. Cell viability was assessed using a resazurin assay at Days 5, 7, and 9. Individual dots represent biological replicates and asterisks (*) indicate statistically significant between group (*p*<0.05).

### CFU formation supports the clonogenic capacity of PE gAD-MSCs

During the CFU assay, gAD-MSCs exhibited a clear temporal progression of colony development. Small colonies emerged by Day 5 and expanded continuously throughout the culture period, resulting in large, densely cellular colonies by Day 14 (Figure 4). The colonies were composed predominantly of spindle-shaped cells and displayed characteristic colony architecture consistent with mesenchymal stem cell cultures. The progressive increase in colony size and cellular density indicates that a proportion of the cultured gAD-MSC population retained clonogenic properties, enabling individual progenitor cells to survive, proliferate, and generate multicellular colonies. These findings provide functional evidence of the clonogenic potential of the isolated gAD-MSCs and complement the viability and morphological characterization results.

**Figure 4.**
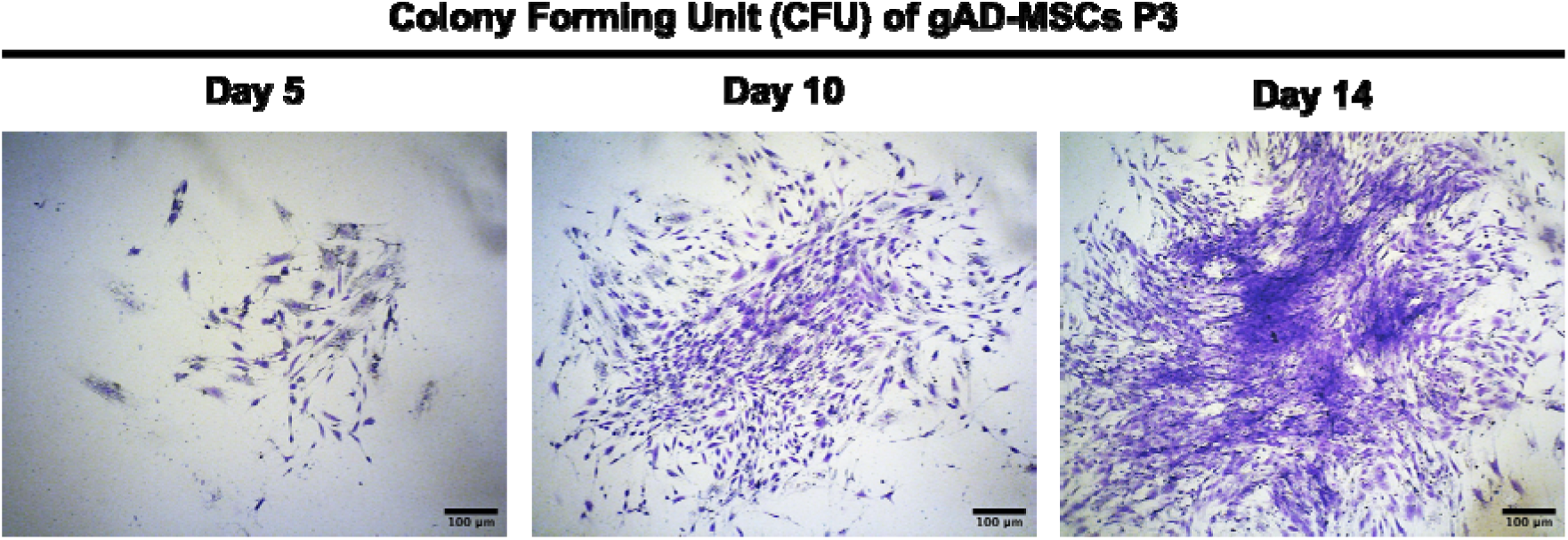
Representative stages of colony development during the colony-forming unit (CFU) assay of gAD-MSCs. Crystal violet-stained colonies were observed throughout the 14-day culture period following low-density seeding. Early colony formation on Day 5, characterized by small clusters of adherent spindle-shaped cells. Progressive colony expansion on Day 10, showing increased cell density and enlargement of the colony area. Mature colony on Day 14, exhibiting a densely packed population of fibroblast-like cells arranged in characteristic swirling patterns. These observations demonstrate the clonogenic capacity of gAD-MSCs and their ability to survive, proliferate, and generate colonies from clonogenic progenitor cells. Scale bars = 100 μm

Quantitative assessment of the CFU assay further demonstrated the colony-forming capacity of gAD-MSCs passage 3. Following 14 days of culture, colonies containing ≥50 cells were counted according to the predefined CFU criterion, yielding an average of 52.0 ± 4.1 colonies (Figure 5). The relatively low variation among replicates indicates consistent colony-forming activity across the cultures. Together with the progressive colony development observed by Crystal Violet staining, the quantitative findings demonstrate that the expanded gAD-MSC passage 3 population retained clonogenic and proliferative capacity, supporting its suitability for subsequent phenotypic characterization and multilineage differentiation analyses.

**Figure 5.**
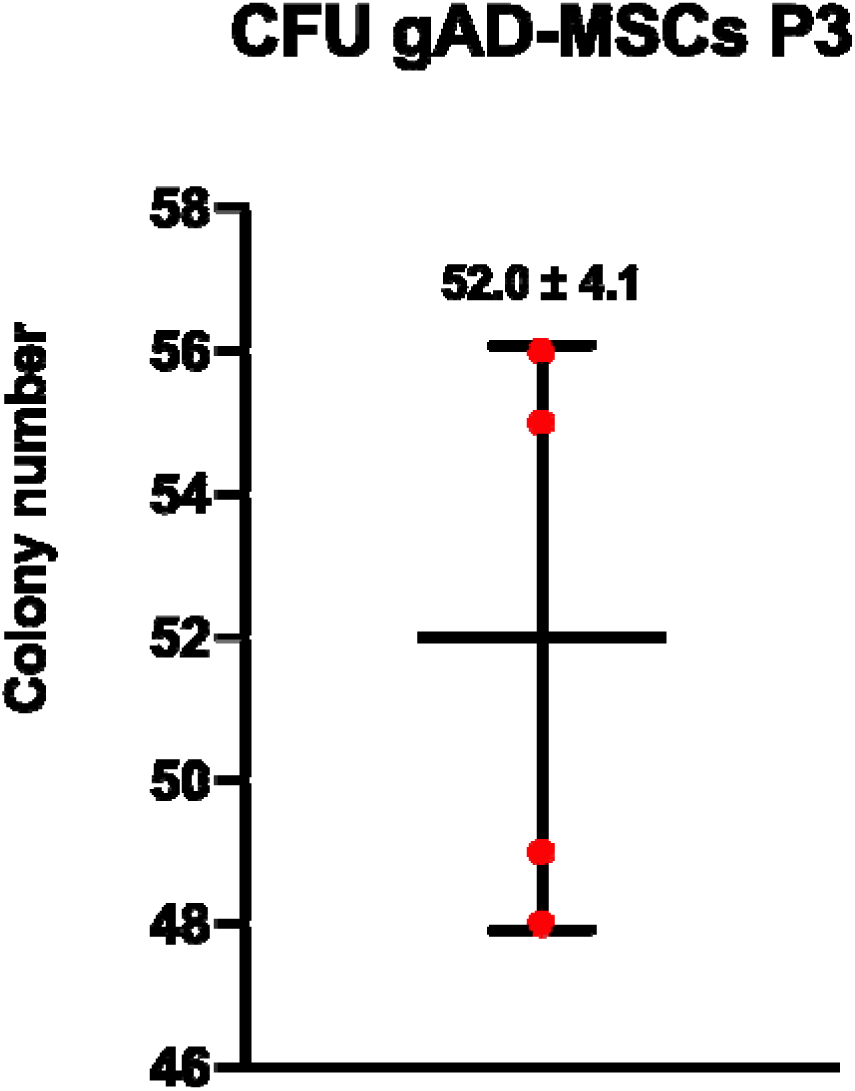
Quantitative assessment of CFU formation by passage 3 gAD-MSCs after 14 days of culture. Following low-density seeding, colonies were visualized by crystal violet staining and counted based on the predefined criterion of ≥50 cells per colony. Individual dots represent biological replicates, and data are presented as mean ± SD.

## DISCUSSION

The present study obtained an adherent cell population from adipose tissue collected at the base of the tail of Peranakan Ettawa (PE) goats via explant culture. Cell outgrowth from the explants was evident by Day 5 and became progressively more extensive during continued culture, with cells exhibiting an elongated, spindle-shaped morphology. These features are consistent with the classical morphological characteristics of mesenchymal stem cells (MSCs), which are generally characterized by adherence to tissue culture plastics and a fibroblast-like appearance under standard culture conditions (1, 6, 7). Importantly, recent studies specifically characterize goat adipose tissue-derived MSCs reported well-spread, elongated cells with clear nuclei, strong adherence, and homogenous morphology under supportive culture conditions, closely resembling the morphological characteristics observed in the present study (6). Similar morphological characteristics have been reported across several veterinary species (8–16). The consistency of these characteristics across different species represents a conserved feature of adipose-derived mesenchymal stem cell populations rather than a characteristic restricted to a particular species. The progressive migration of cells from the explant and their subsequent local expansion indicate the emergence and expansion of adherent adipose-derived mesenchymal stem cells. Nevertheless, morphology and plastic adherence alone cannot definitively establish MSC properties, these characteristics should be interpreted as initial components of MSC characterization rather than as evidence of stemness per se.

Following serial expansion, gAD-MSCs passage 3 retained a consistent spindle-shaped morphology and adherence to the culture surface, with relatively limited morphological heterogeneity. This pattern is consistent with observations in sheep, in which adipose -and bone marrow-derived MSCs exhibited more heterogeneous morphology during early passages at later passage, including P3 (17, 18). This morphological stability is important for establishing a reproducible cell population for subsequent functional characterization. The recent comprehensive characterization of gAD-MSCs similarly demonstrated that appropriate culture conditions could maintain fibroblast-like morphology, adherence, metabolic activity, clonogenicity, and proliferation, although these properties were influenced by basal culture medium (6). Thus, the relatively homogenous morphology observed in the present P3 cultures suggests that the standard culture condition used were compatible with the maintenance and expansion of the adherent gAD-MSCs population. However, it is more appropriate to describe this finding as morphological consistency or enrichment of an adherent mesenchymal population, rather than claiming that serial passage itself eliminated all non-MSCs phenotypes. In accordance with current ISCT nomenclature recommendations, the designation MSCs should ideally be supported by a combination of tissue origin, immunophenotypic characteristics, and relevant functional assays rather than morphology alone (7). The subsequent phenotypic and multilineage differentiation analysis in the present study is therefore important for establishing the identity and functional properties of the P3 population.

The resazurin assay demonstrated a progressive increase in signal following seeding of 2 × 10^3^ gAD-MSCs P3, with a statistically significant increase between Day 3 and Day 5. This pattern is consistent with an increase in the metabolically active cell population during culture and is consistent with the previous observation that gAD-MSCs exhibit measurable metabolic activity and proliferative expansion under appropriate culture conditions (6). Resazurin reduction primarily reports cellular metabolic activity and is not a direct measurement of cell number, proliferation, or cell death. Changes in cellular metabolic state can alter the assay signal independently of changes in cell number, and the assay signal may also be affected linearly by cell density, incubation time, and resazurin concentration (19). Recent methodological work has consequently emphasized optimization and standardized reporting of resazurin-based assays to improve the reliability of quantitative interpretation (20). Therefore, the significant increase between Day 3 and 5 in the present study is best interpreted as an increase in metabolic activity associated with the expansion of the viable cell population. When considered together with the observed morphological expansion and subsequent CFU findings, the resazurin result report supports the ability of gAD-MSCs P3 to remain metabolically active and expand under the condition use in this study.

The CFU assay provided complementary functional evidence that the proportion of the gAD-MSC P3 population retained the ability to generate colonies following low-density initial culture. Small colonies were detectable during the early phase of the assay and progressively increased in size and cellular density until Day 14. The development of discrete behavior expected of MSC-containing mesenchymal populations provides information distinct from that assessed by resazurin. This interpretation is also supported by recent work in gAD-MSCs, in which colony-forming assays were used to evaluate clonogenic capacity (6). The relatively low variation in colony number among the present replicates further suggests reproducible colony-forming activity in the P3 population. Importantly, CFU provides information that is distinct from the metabolic activity assessed by resazurin, as it evaluates the ability of a subset of cells to survive under low-density conditions and generate multicellular colonies. Accordingly, the combined morphology, metabolic activity, and CFU results establish a strong preliminary functional profile of the expanded gAD-MSCs population derived from PE goats, while the subsequent analysis are necessary to provide a more comprehensive characterization.

Taken together, adipose-derived cells from Peranakan Ettawa (PE) goats exhibited characteristic MSC-like morphology, sustained metabolic activity, and clonogenic capacity following expansion to Passage 3. These findings provide a preliminary characterization of gAD-MSCs and establish a baseline cellular profile for further phenotypic and multilineage differentiation studies. Collectively, the results support the potential of PE goat-derived MSCs as a locally relevant cell source for further development in veterinary regenerative medicine.

## AUTHOR‘S CONTRIBUTION

TB contributed to conceptualization, supervision, methodology, resources, project administration, and manuscript review and editing. IP contributed to conceptualization, methodology, investigation, data curation, formal analysis, visualization, writing of the original draft, and manuscript review and editing, and served as the corresponding author. MP and VB contributed to the investigation, methodology, resources, validation, and manuscript review and editing. NP contributed to supervision, formal analysis, and manuscript review and editing. All authors read and approved the final version of the manuscript.

## SUPPLEMENTARY MATERIALS

There is no supplementary material.

## DATA AVAILABILITY

The datasets generated and or analyzed during the current study are available from the corresponding authorupon reasonable request.

## ETHICS APPROVAL

All animal procedures were conducted in accordance with the guidelines of the Animal Ethics Committee of the Faculty of Veterinary Medicine, Universitas Gadjah Mada, and were approved under protocol number 03/EC-FKH/Int./2026.

## DECLARATION OF GENERATIVE AI

During the preparation of this manuscript, the authors used ChatGPT (OpenAI) to assist with language editing and improvement of English grammar and readability. No generative AI was used to generate scientific data, analyze data, or draw scientific conclusions. The authors reviewed and edited the output and take full responsibility for the manuscript’s content.

