## Supplementary figures and images for "Morphological and phenotypic characterization of adipose-derived mesenchymal stem cells isolated from locally adapted Indonesian goat breed"

### Figure file

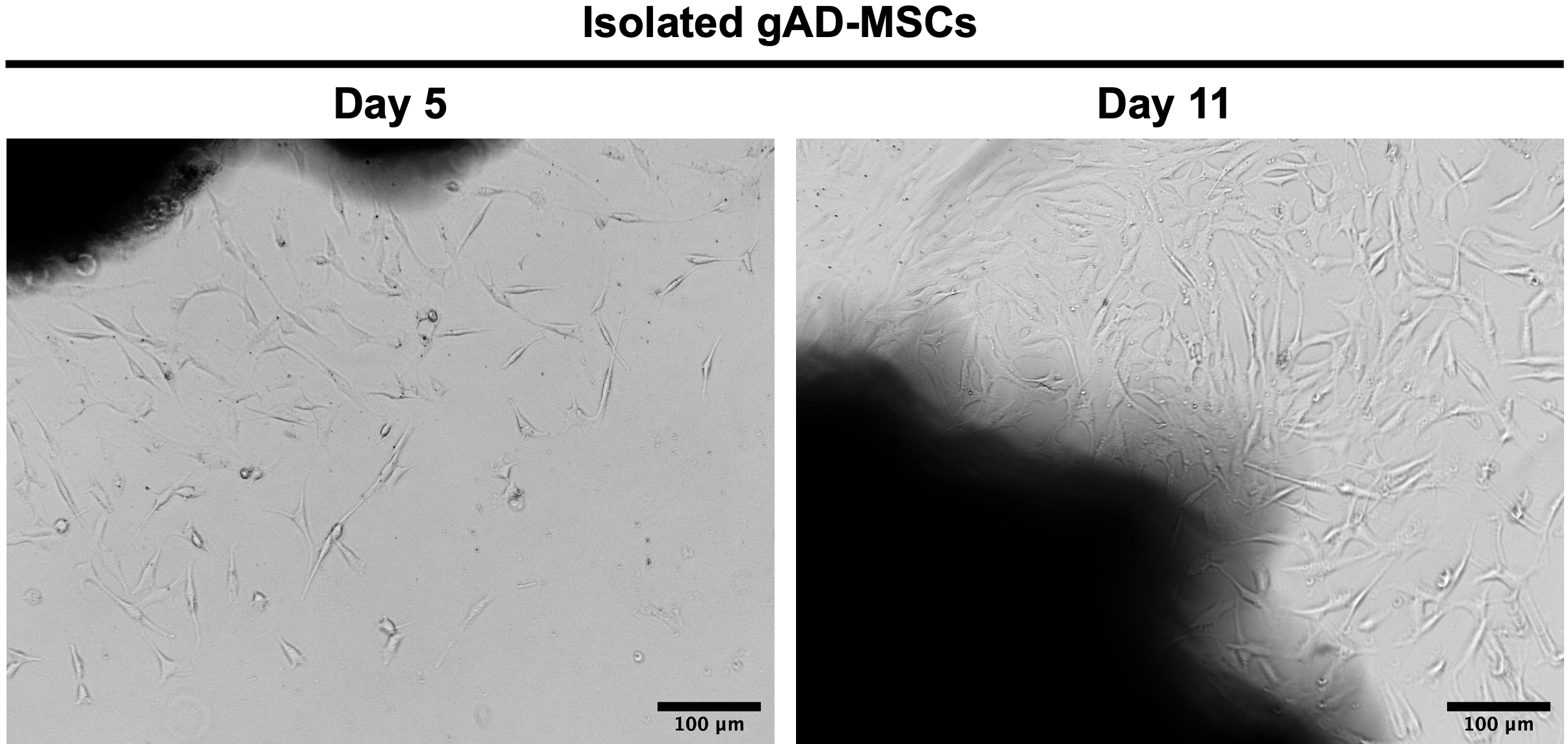

Figure 1

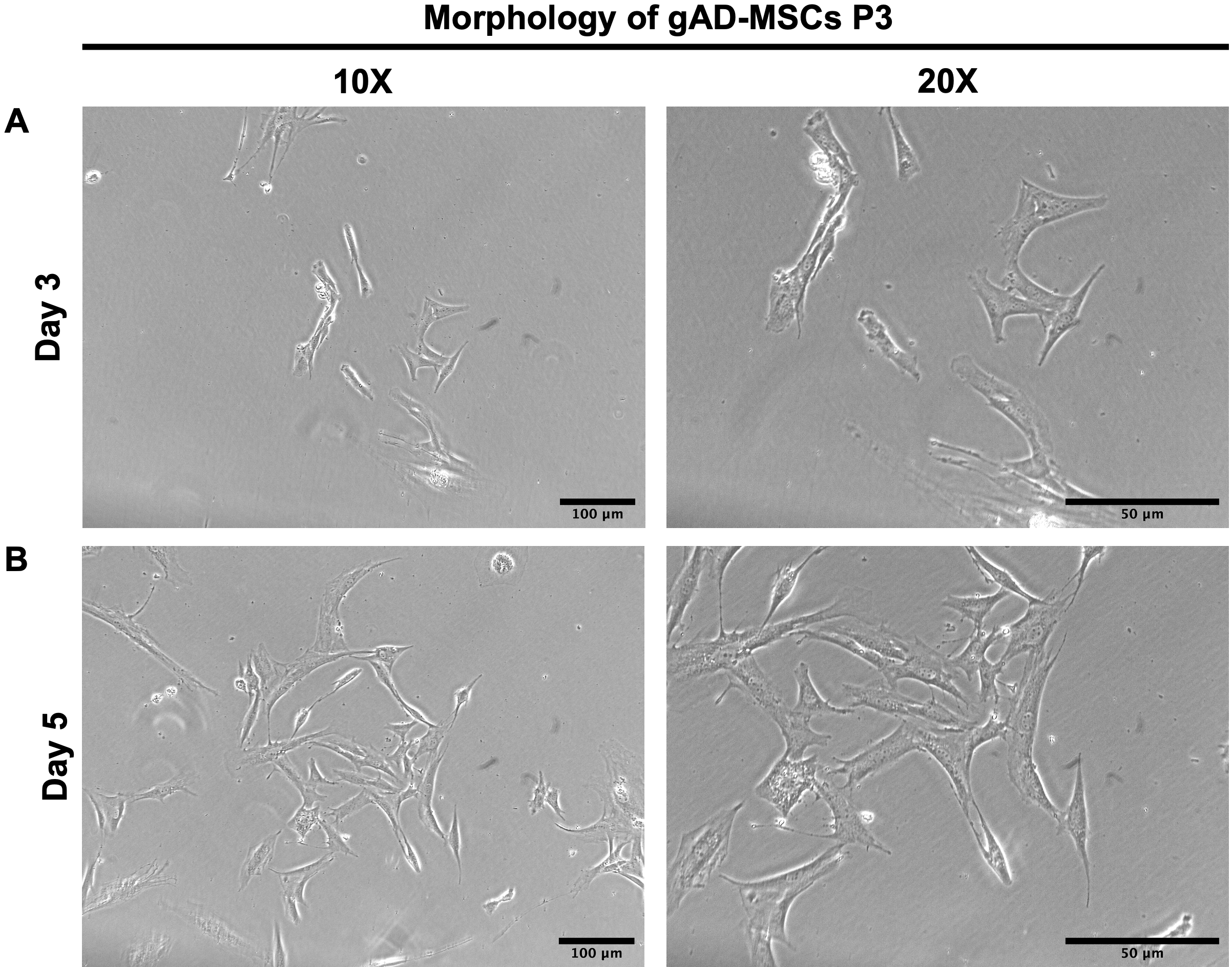

Figure 2

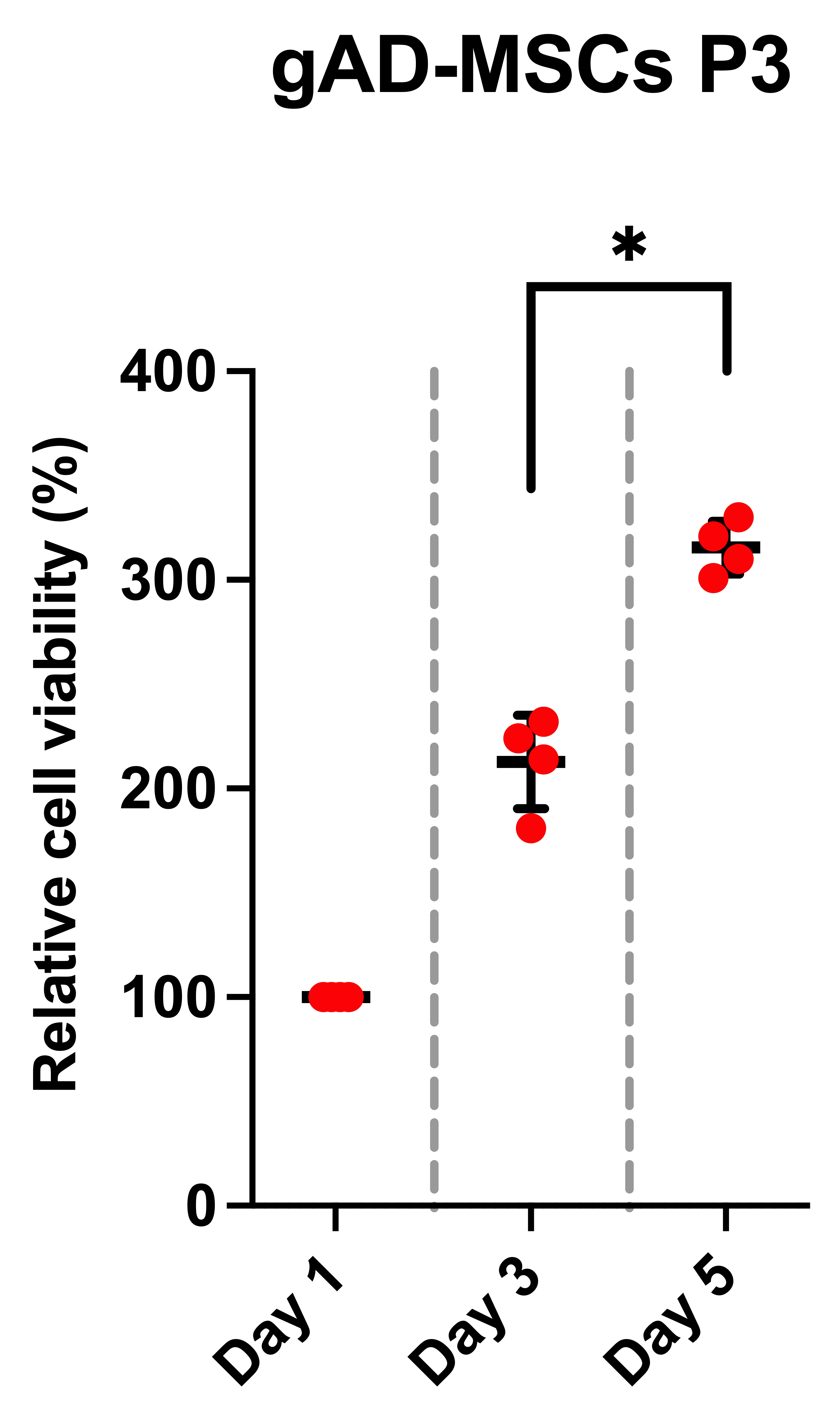

Figure 3

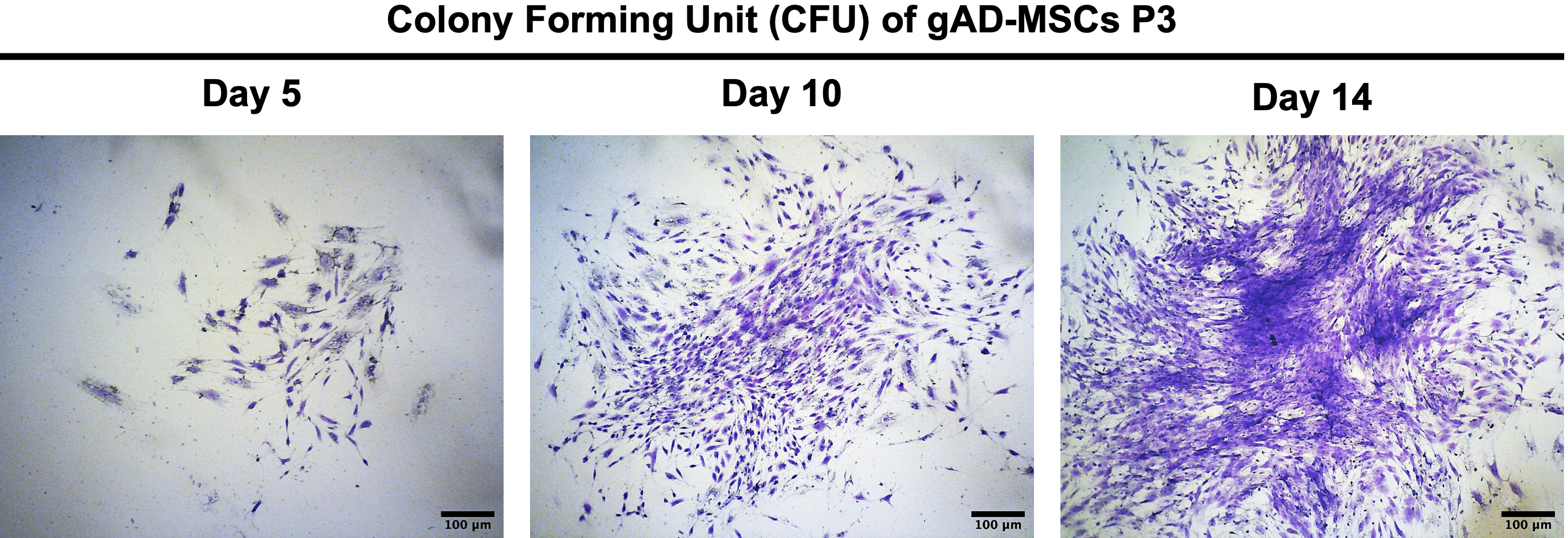

Figure 4

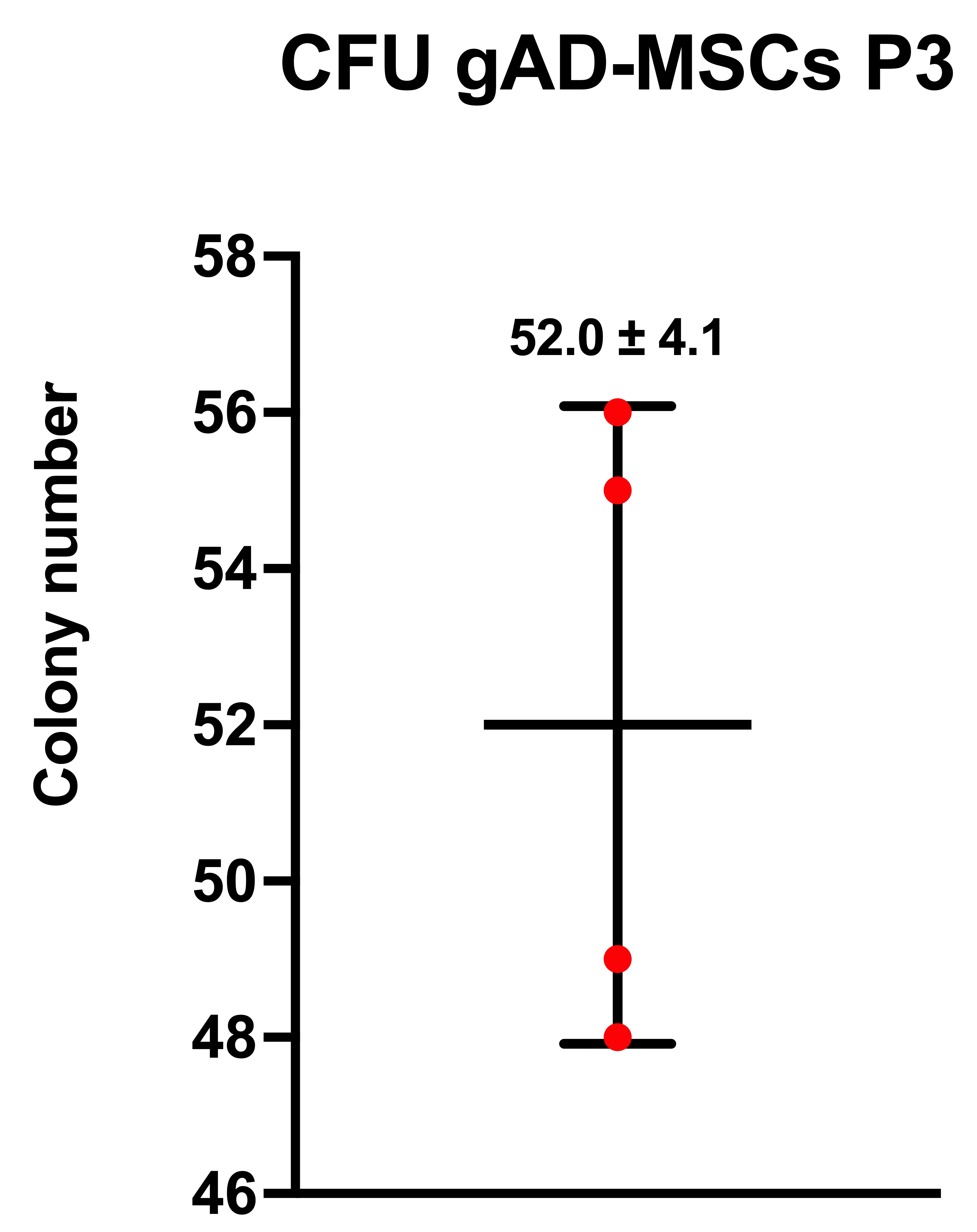

Figure 5
